# Influence of different landscape configurations on insect pollinators of important horticultural crops in a western Himalayan landscape

**DOI:** 10.1101/2023.12.23.573179

**Authors:** Susmita Khan, Bitapi Sinha, V.P. Uniyal

## Abstract

Pollinators play critical roles in ecosystem stability and agricultural productivity, and their decline has become a global concern. One of the main reasons for pollinators decline is changing landscape configuration. We conducted this study in Kullu Valley, Himachal Pradesh, India to evaluate varied landscape configuration impacts on the diversity, richness, and abundance of insect pollinators in apple and plum orchards. Five different landscape configurations were considered for the study. The survey spanned 28 orchards across the valley during the blooming periods of 2021 and 2022. Standard sampling protocols were used to collect the pollinator specimen. We recorded a total of 107 insect species across 67 genera, 29 families and five orders (Hymenoptera, Lepidoptera, Diptera, Coleoptera, and Thysanoptera) from different orchards. Hymenoptera and Diptera were the most abundant pollinators in the study area. Pollinator richness and diversity were higher in orchards located within 1-km of natural forests and in lower elevations (>1500 meters). Orchards near large settlements, agricultural lands and complete modernised management practice showed comparative less pollinator diversity and abundance. This study demonstrates the influence of diverse land-use practices on pollinators on a landscape scale and provides insights for developing effective conservation strategies in the Himalaya.

## Introduction

Pollinators play a crucial role in ecosystems by maintaining biodiversity, sustaining natural habitats, and supporting agricultural productivity (Misganaw et al. 2017). Beyond their ecological significance, pollinators contribute immensely to global food production, ensuring the growth of many crops for human and animal sustenance (Klein et al. 2007). Their economic contribution to global agriculture has been estimated at 153 billion pounds (€) annually (Gallai et al. 2009). Apart from bees, there are various non-bee pollinators such as flies, wasps, moths, butterflies, beetles, ants, birds, and bats, who play a crucial role in pollination by providing 39% of crop pollination visits, however, their contribution is little explored (Rader et al. 2016, 2020). Studies have recorded that 84% of total pollen is carried by non-syrphid dipteran flies (Orford et al. 2015).

In recent years, there has been a significant decline in both the abundance and diversity of insect pollinators (Ghazoul 2005; Steffan-Dewenter et al. 2005; Biesmeijer et al. 2006; Williams and Osborne 2009). While much attention has been given to the decline of managed honeybee populations due to Colony Collapse Disorder (Oldroyd 2007), several wild bee species have also experienced sharp declines and, in many cases, have disappeared from their historic natural ranges (Biesmeijer et al. 2006; National Research Council 2007; Potts et al. 2010; Cameron et al. 2011)

In the past few years, a disturbance in crop pollination services due to a decline in the number and diversity of pollinators has been noticed throughout the Hindu Kush Himalayan region (Partap and Partap, 1997, 2002; Partap et al. 2001; Ahmad et al. 2003; Partap 2010a, 2010b). Studies proved that changes in Landscape configuration, including alteration of natural habitats for anthropogenic use and agricultural intensification, have been identified as one of the major factors of pollinator declines (Kennedy et al. 2013; Vanbergen et al. 2013). Other key factors causing pollinator decline include an increase in monoculture-dominated agriculture and the use of pesticides (Verma and Partap 1993; Aizen and Feinsinger 1994; Partap and Partap 1997, 2002; Allen-Wardell et al. 1998; Ahmad et al. 2003; Partap 2010 a, 2010b).

Forests play a vital role by offering habitats for nesting, hibernation, and food resources for diverse pollinator species. Research indicates that orchards near forests host a higher diversity of insect pollinators compared to those situated farther away (Sharma and Gupta 2010; Ulyshen et al. 2023). Thus, any reduction in forested areas, whether due to natural disasters or anthropogenic interventions, adversely affects pollinator abundance (IPBES 2016).

Climate change can impact pollinators by affecting the lifecycle of natural pollinators due to altered local weather conditions (Partap and Partap 2001, 2002). Identifying the influence of landscape changes on pollinator communities is critical to prevent additional pollinator loss and to develop effective strategies for their protection in human-influenced landscapes (Viana et al. 2012).

Himachal Pradesh stands out as a prominent horticultural region in India, characterized by diverse agro-climatic zones spanning from subtropical areas to high-altitude cold deserts. This variety allows for the successful cultivation of a broad spectrum of horticultural crops. Consequently, there has been a notable shift in the agricultural landscape, witnessing a significant transition from traditional farming practices to the cultivation of high-value horticultural crops (Sharma and Chauhan 2013; Slariya 2014). The commonly grown temperate fruit crops are apple, plum, peach, cherry, pear, and apricot which occupy approximately 35 percent of the total area of the hill state (Sharma and Rana 2015). There have been past studies on different insect pollinators in Himachal Pradesh including the Kullu district (Sharma and Gupta 2010; Sharma and Mitra 2011; Pratap et al. 2012; Thakur and Mattu 2014; Mattu and Bhagat 2015; Sharma et. al. 2015; Kapkoti et. al. 2016; Sharma et. al. 2016; Chauhan et. al. 2021; Arya and Badoni 2023). However, studies on pollinator diversity of important fruit crops in different land-use categories are still lacking.

Therefore, the present study was conducted to know the effect of different landscape categories on the diversity, richness, and abundance of insect pollinators of apple and plum crops in the Kullu Valley of Himachal Pradesh.

## Materials and Methods

### Study area

The survey was conducted within the Kullu Valley situated in Himachal Pradesh, covering a stretch of 76 km along the Beas River between 32°21’13.90”N and 77° 7’38.57”E to 31°43’27.64”N and 77°12’52.54”E. The valley stretches to a width of 3-4 km on both sides. Elevations of the valley range between 800-3000 meters and experiences an average annual rainfall of 8000mm, with temperatures varying between 0°-38°C. The region has been significantly impacted by tourism activities (Gardner et al. 2002) and also faced rapid land-use changes over the past years (Vaidya et al. 2018; Negi and Irfan 2022). According to data of HP State Council for Environment Science and Technology, Kullu district falls under a very high vulnerability zone due to various natural and anthropogenic hazards which makes this land highly fragile and faces frequent alterations in the land-use category(https://hpsdma.nic.in) (Figure 1). Horticultural practices in this area primarily involve mixed cropping, with key commercial fruit crops including Apple, Plum, Pear, Pomegranate, Kiwi, and Peach (Verma et al. 2010). The survey was conducted in 28 plum and apple orchards across the entire valley during the full blooming period of the orchards in the years 2021 and 2022 (Figure 2).

**Figure 1.**
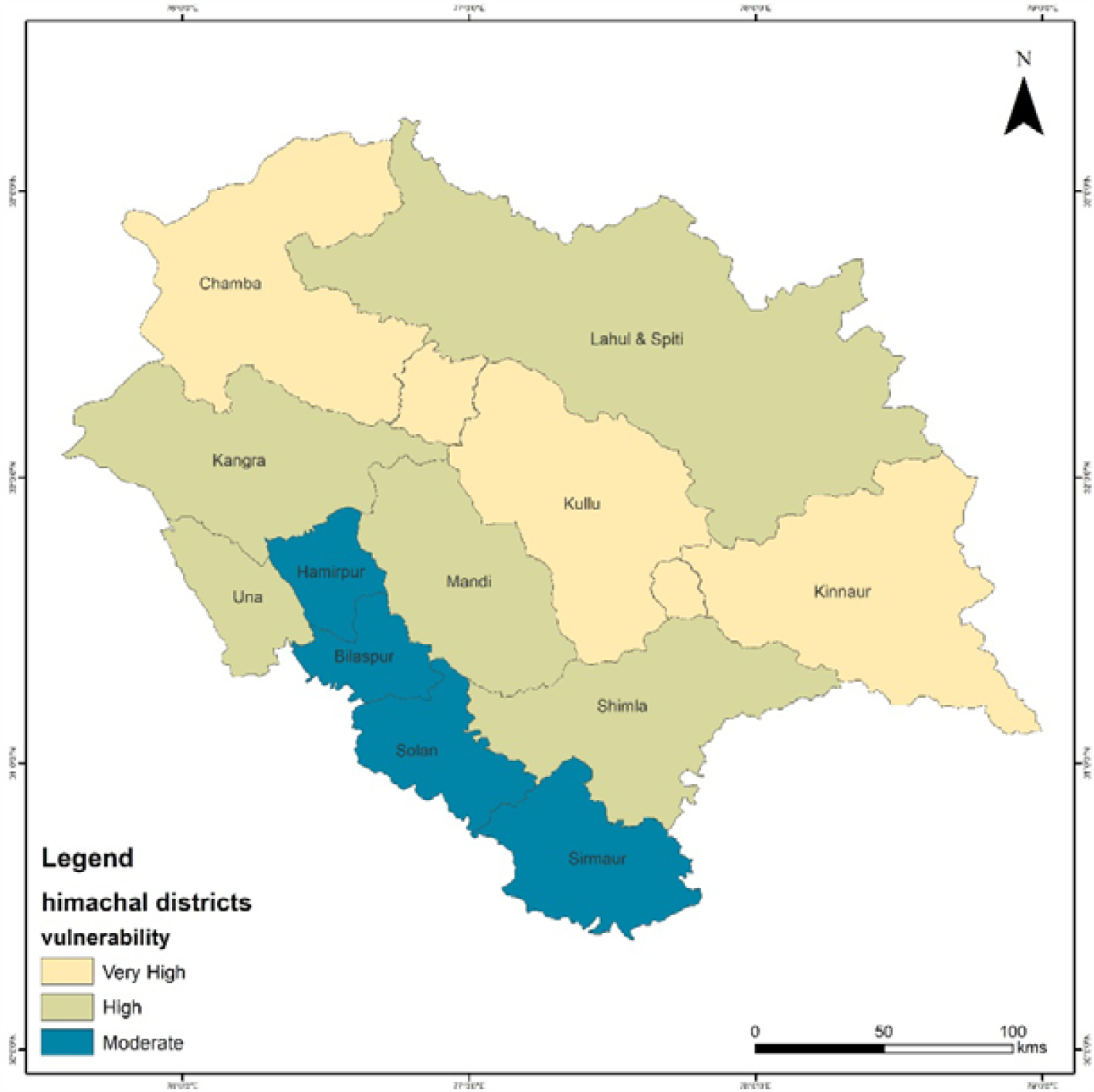
Landscape vulnerability map of Himachal Pradesh (Adopted from: Himachal Pradesh state disaster management authority)

**Figure 2.**
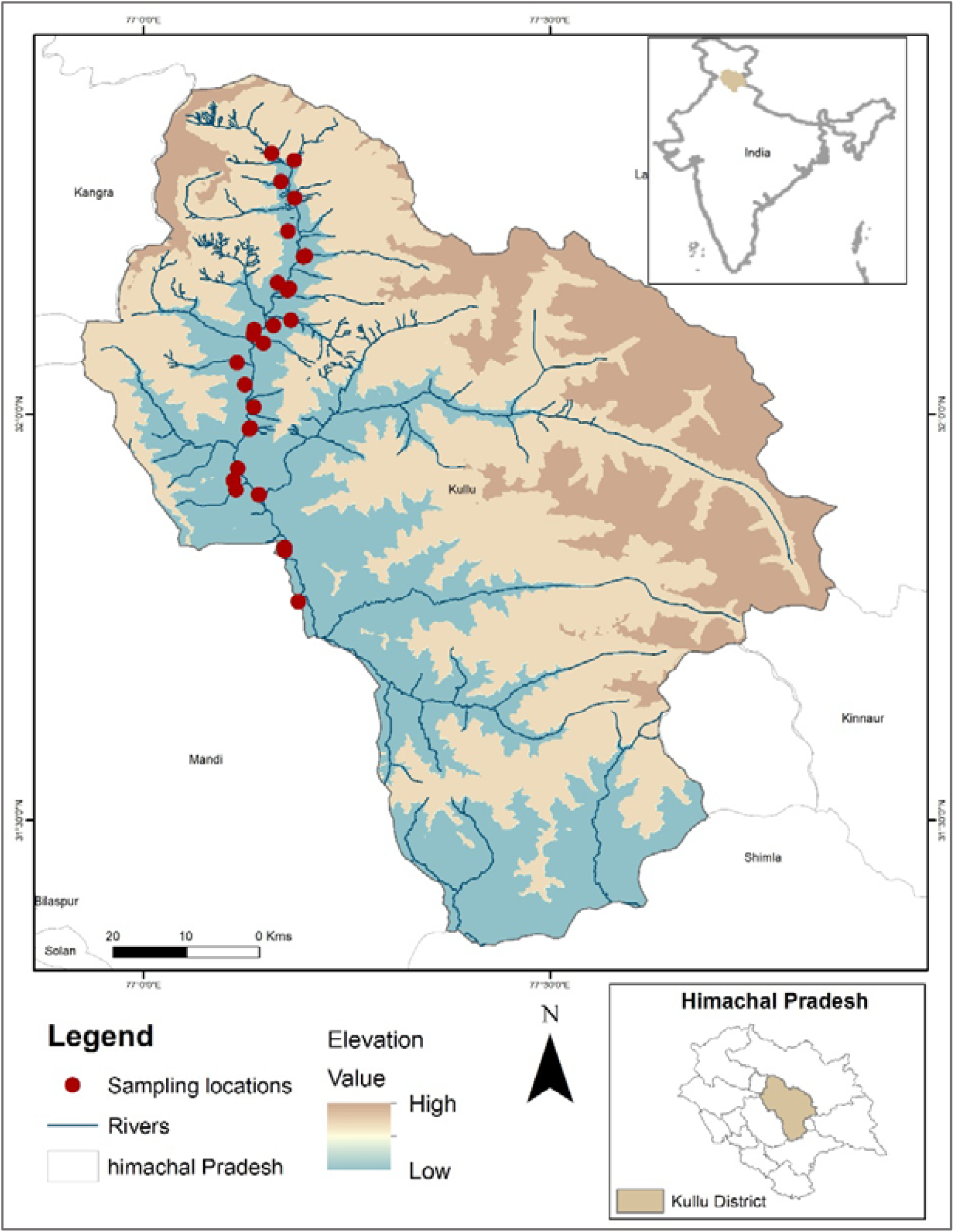
Map of study area

The study aimed to assess the influence of various landscape configurations on pollinator diversity and richness. Sampling locations were chosen based on five landscape variables; i.e. proximity to large forest patches (i.e. less than 1km, 1-2 km, more than 2kms), elevation gradients (i.e. below 1500m, 1500-2000m, and above 2000m), the presence of neighboring agricultural fields, the proximity of large settlements, and the types of management practices employed (e.g. modern, modern and traditional).

### Methodology

Sample collection was done within a 50m x 50m plot during daylight hours from 9 am to 5 pm in each location. Both active methods such as hand picking, sweep netting, and aerial netting (Joseph 1990; Arora 1990) and passive methods like pan traps were employed to collect insect samples. The pan traps, colored in UV-bright yellow, white, and blue colours (Westphal et al. 2008) and filled with soapy water, were arranged in clusters, and placed 5 meters apart from each other. Focal observations (Gibson et al. 2011) were conducted for 10-minute within 1m x 1m plots along flowering branches at three different time frames: 09:30-10:30 h, 12:30-1:30 h, and 15:30-16:30h. All specimens collected using wet methods were washed with plain water and preserved in 75% ethanol. Those obtained through active methods were euthanized using ethyl acetate in a killing jar and preserved in dry pouches. In the laboratory, specimens underwent further processing, including washing, drying, pinning, and labelling, followed by identification using standard identification keys.

The samples gathered through diverse sampling methods were combined for the analysis of species diversity and richness. To assess the influence of distinct landscape configurations on pollinator diversity, separate analyses were performed for each land use category. Our analysis involved assessing Shannon and Simpson’s diversity indices, the Shannon evenness index, species richness, and Chao-1. We have also run regression analysis (Generalized Additive Model) to see the relation between pollinator abundance with different elevational gradients and changing distance to forests. Data were analyzed in Excel and R studio using biodiversityR (Kindt 2019) and vegan (Dixon 2003) packages. Observations were made on warm, sunny days when pollinators are more active, avoiding rainy and cloudy days to reduce bias.

## Results

In the study area, a total of 107 insect species were observed across 67 genera and 29 families, belonging to five orders: Hymenoptera, Lepidoptera, Diptera, Coleoptera, and Thysanoptera. The order Hymenoptera comprised 40 species within 19 genera and 8 families, while Diptera accounted for 39 species distributed among 23 genera and nine families. Lepidoptera included 18 species across 17 genera and five families, followed by Coleoptera with seven species under seven genera and six families. Lastly, Thysanoptera was represented by three species within one genus and one family (Table 1, Figure 3). The Shannon (Shannon-H) and Simpson’s (Simpson 1-D) diversity index was reported 4.137 and 0.978 respectively. Shannon Evenness index (e^H/S) was calculated 0.581 followed by Chao-1 as 107.1 (Table 1).

**Table 1.**
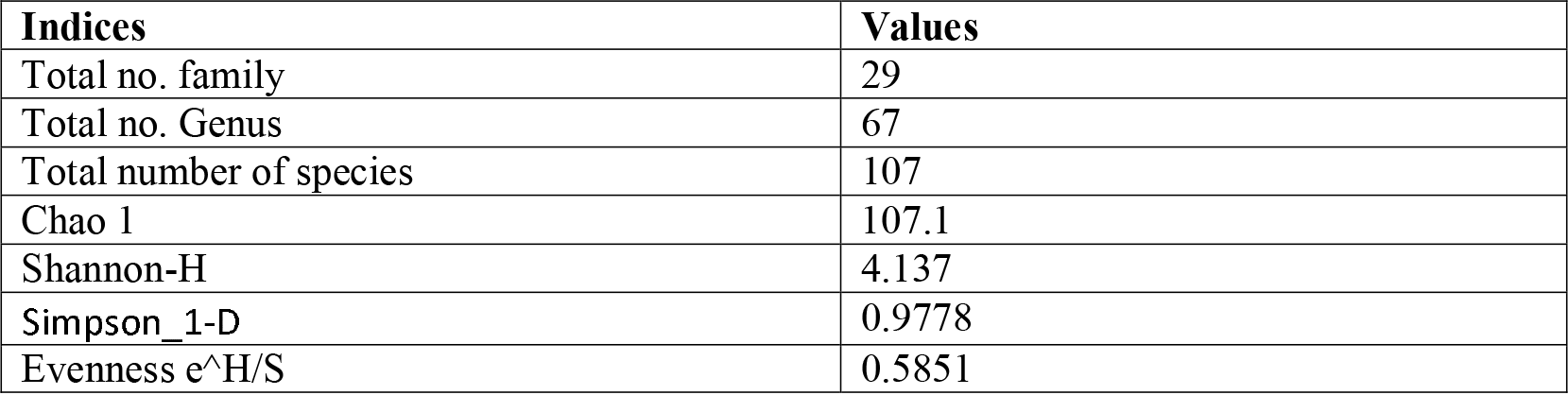
Details of different indices of insect visitors in study area.

**Figure 3.**
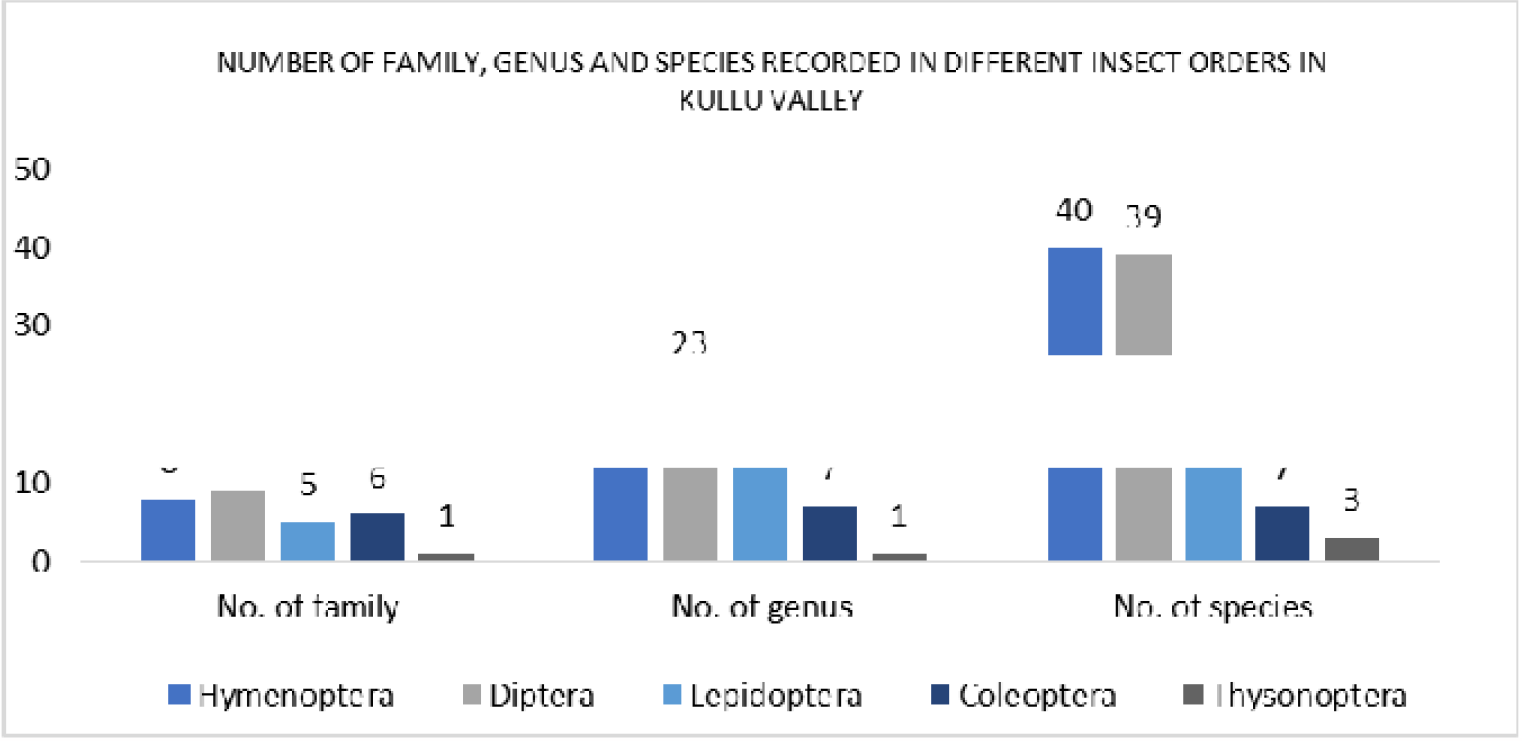
Account of family, genus, and species of different insect orders

To evaluate the impact of various land use categories on pollinators, focus was placed on three insect orders i.e. Hymenoptera, Diptera, and Lepidoptera, considering their higher efficiency as pollinators.

When considering the proximity to forest areas, the orchards located less than one kilometre from natural forests exhibited the highest species richness and diversity for both Hymenoptera and Diptera. This trend was followed by orchards situated within a 1-2 km distance from forests and those situated more than two kilometres away. Concerning Lepidoptera, the highest species richness was observed in orchards within 1-2 km from forests, followed by orchards within one km and those far than two kms from forests. However, the maximum species diversity was recorded in orchards less than one km from natural forests, followed by orchards within a 1-2 km range and those situated more than two km away (Figure 4). The abundance of insects exhibited a similar pattern across all three insect orders, with a decrease in abundance as the distance from the forest increased (Figure 5).

**Figure 4.**
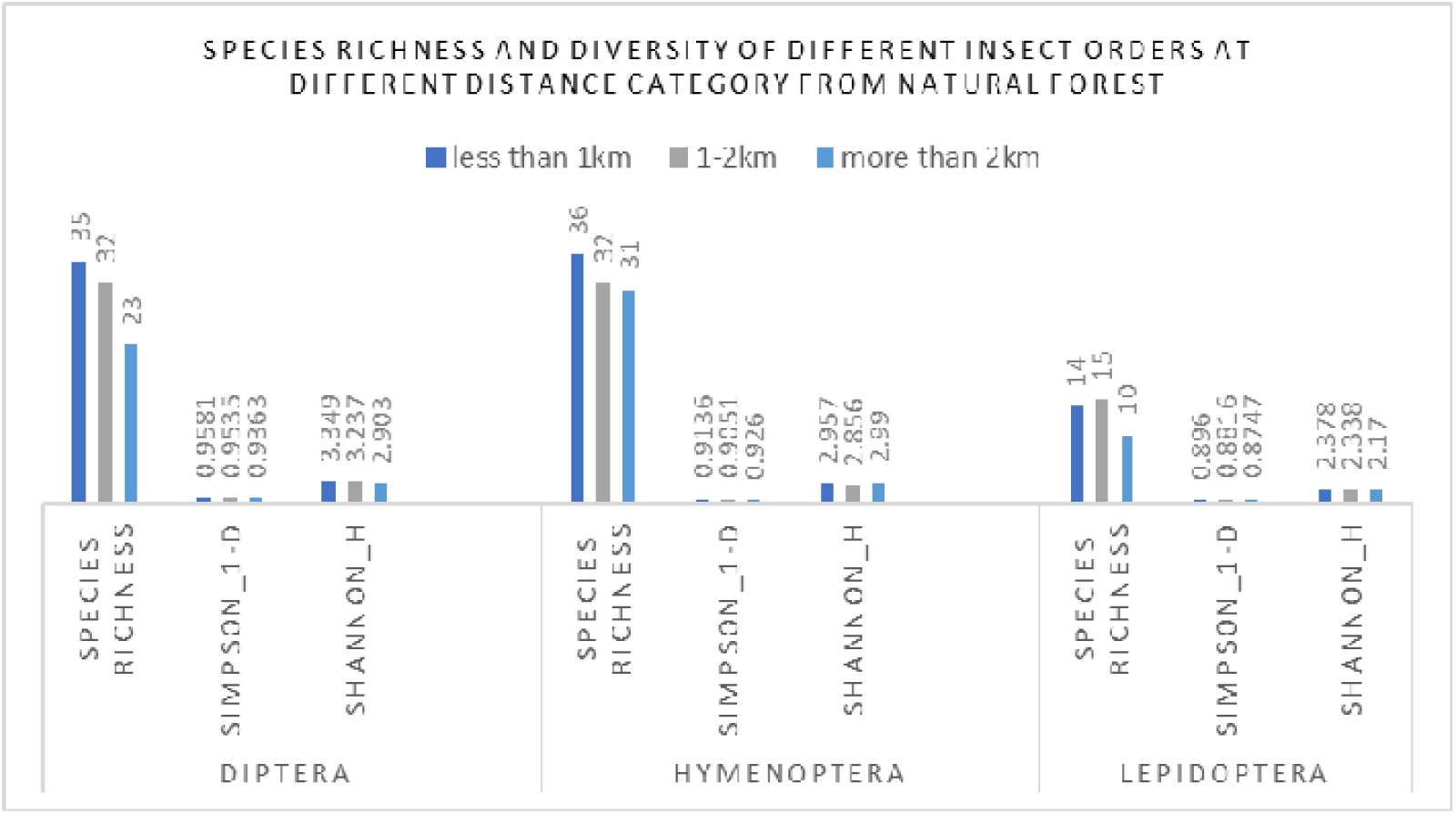
Account of species richness and diversity with changing distance from forest

**Figure 5.**
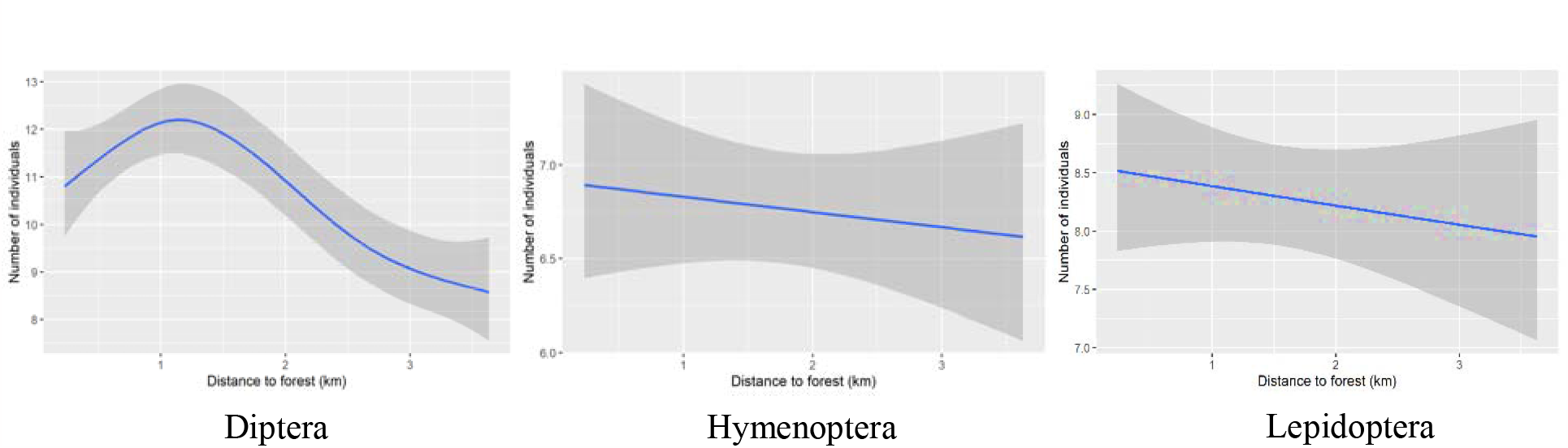
Relation between pollinator abundance and distance to forest

### The influence of elevation on pollinator diversity

The highest richness and diversity of pollinators were observed in orchards situated below 1500 meters above sea level, followed by orchards within the 1500m-2000m elevational range. The lowest richness and diversity were found in orchards located above 2000 meters elevation for both Hymenoptera and Lepidoptera. However, Diptera exhibited slightly different results, showing the highest richness and diversity in the mid-elevation band of 1500m-2000m, followed by orchards below 1500m, and the lowest in orchards above 2000 metres (Figure 6). Hymenoptera and Diptera displayed increased abundance with rising elevation. However, no significant patterns in abundance were observed for Lepidoptera across different elevational bands (Figure 7).

**Figure 6.**
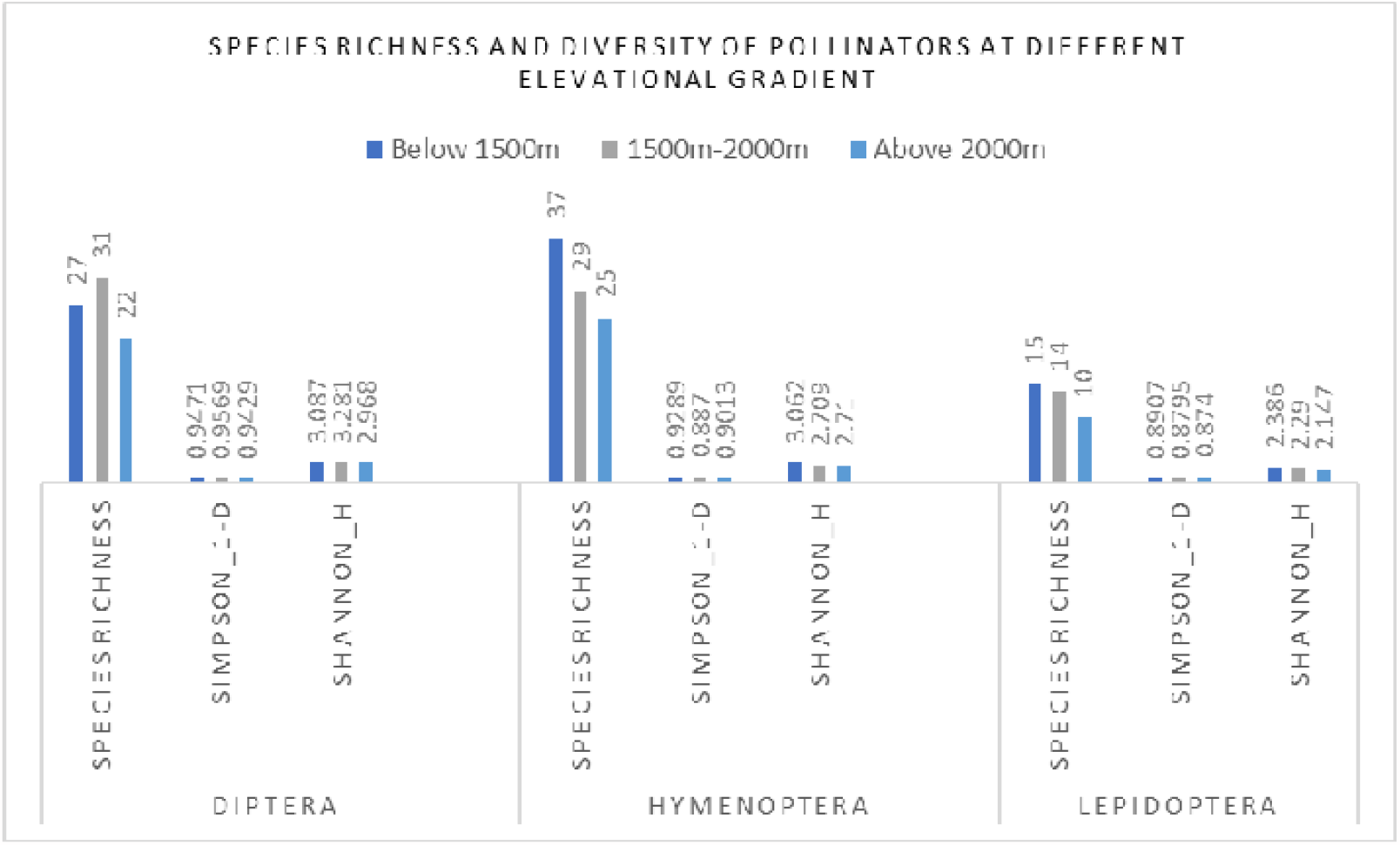
Account of species richness and diversity with elevational gradients

**Figure 7.**
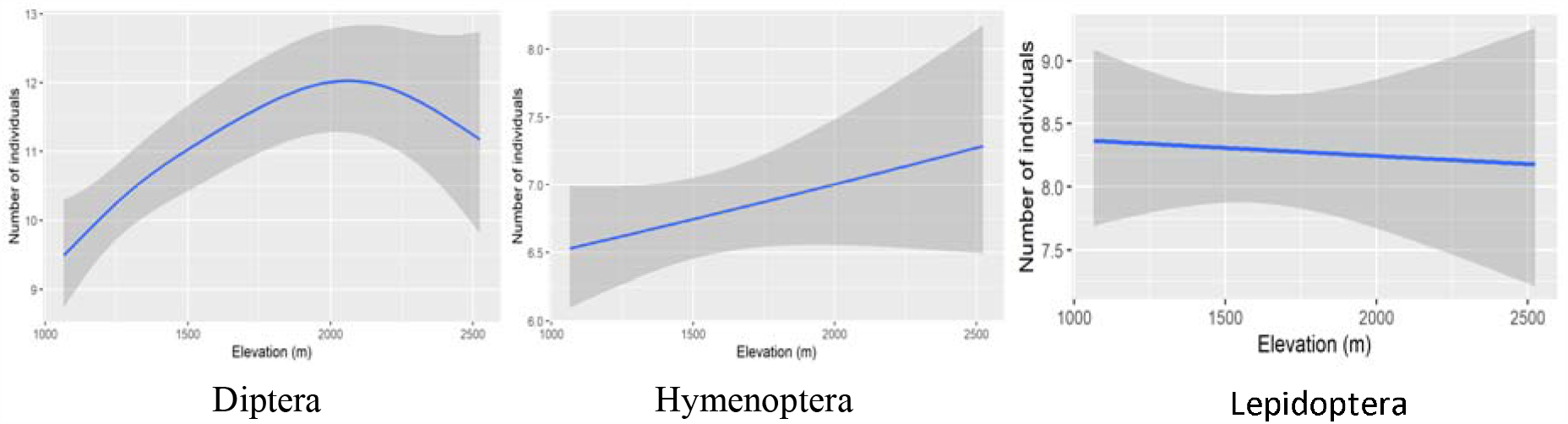
Relation between pollinator abundance and elevational gradient

Presence of agriculture near orchard bed showed higher diversity and richness of dipteran insects where the opposite in case of orders hymenoptera and lepidoptera were observed, where higher insect diversity and richness were found in orchards which is not associated with agricultural field nearby (Figure 8). Most abundant species recorded in orchards present near agricultural land included *Apis cerena indica* (25%), *Apis mellifera* (16%), *Megachilae lanata* (6.3%), *Bombus haemorrhoidalis* (5.1%), and *Amegilla zonata* (4.2%) from the order hymenoptera; *Chrysoma megacephala* (8.81%), *Eristalis tenax* (7.82%), *Eristalis similis* (5.76%), *Eupeodes latifasciatus* (5.76%), *Eupeodes corollae* (5.31%), *Sphaerophoria Indiana* (5.17%) from the order Diptera.

**Figure 8.**
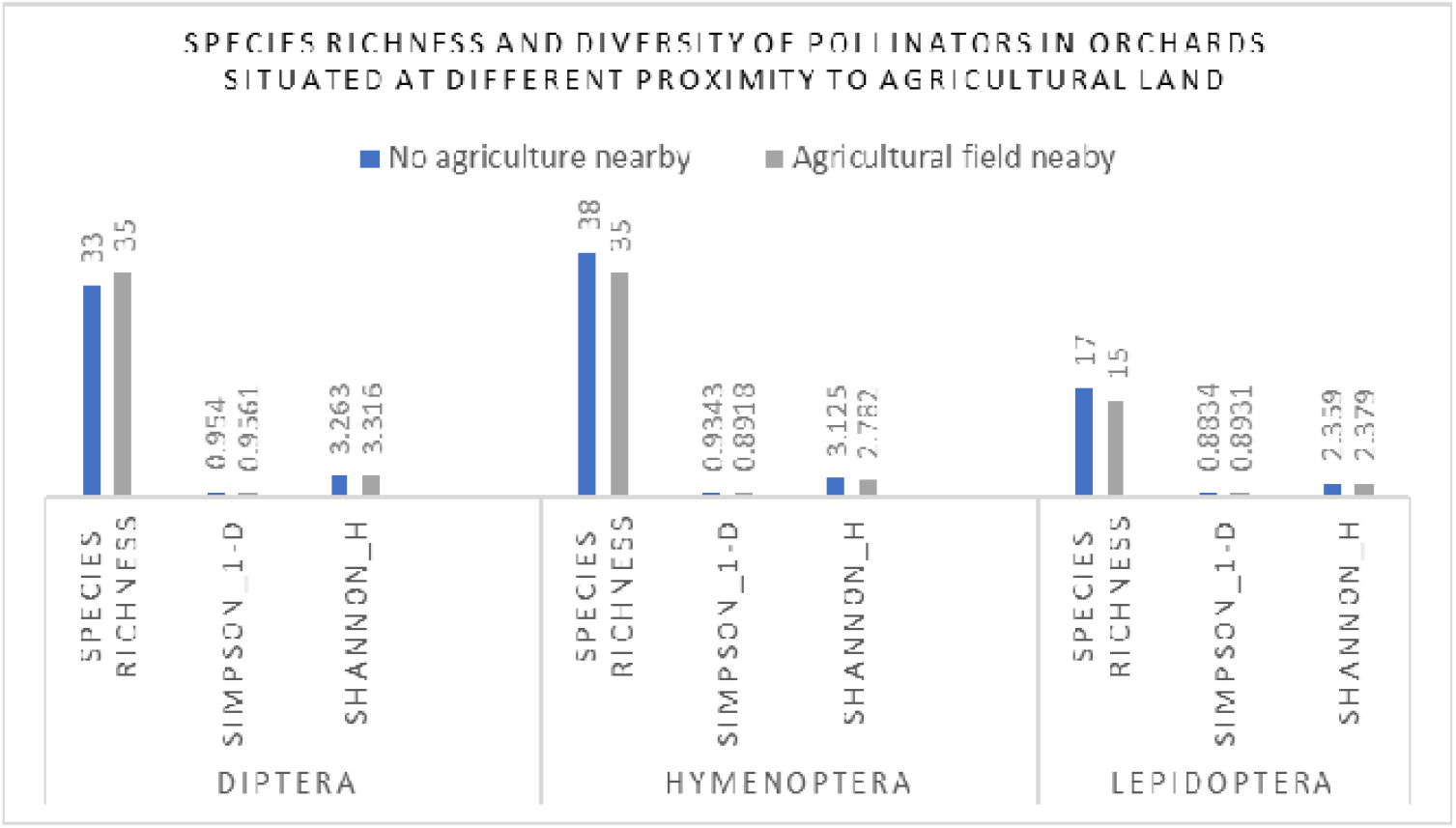
Account of species richness and diversity in orchards at different proximity to agricultural lands

On the contrary, most abundant species recorded from the orchards that does not have agricultural land nearby include *Apis cerena indica* (17.7%), *Apis mellifera* (9.2%), *Megachilae lanata* (7.86%), *Campsomeris sauteri* (5.4%), *Bombus haemorrhoidalis* (4.43%), *Amegilla zonata* (4.33%), *Ceratina smaragdula* (4.12%) and *Xylocopa latipes* (3.82%) from the order hymenoptera; *Eristalis tenax* (9.65%), *Meliscaeva cinctella* (8.32%), *Eupeodes corollae* (7.22%), *Episyrphus balteatus* (5.72%) and *Chrysoma megacephala* (5.41%) From the order Diptera.

In case of lepidoptera most abundant species included, *Aglais caschmirensis* (19.4% & 14.54%), *Colias fieldii fieldii* (16% & 16.34%), *Pieris brassicae* (14.71% & 12.6%), *Eurema hecabe* (10.15% & 10.19%), *Pieris canidia* (7.68% & 10.94%) and *Kaniska canace* (7.68% & 8.9%) in orchards away from agricultural land and orchards near to agricultural land respectively.

Significantly higher insect richness and diversity were observed in the Hymenoptera and Diptera orders in orchards that implemented both modern and traditional management practices, whereas no notable difference was observed in the case of Lepidoptera (Figure 9). In orchards employing both modern and traditional management methodologies, the most abundant species in the Hymenoptera order included *Apis cerena indica* (19.62%), *Apis mellifera* (13.66%), *Megachilae lanata* (7.03%), *Osmia caerulescence* (5%), *Amegilla zonata* (4.73%), *Bombus haemorrhoidalis* (4.73%), Campsomeris sauteri (4.73%), and *Ceratina smaragdula* (4.66%). For the Diptera order, prevalent species were *Eristalis tenax* (8.71%), *Chrysoma megacephala* (7.7%), *Episyrphys balteatus* (6.49%), *Eristalis similis* (5.74%), and *Eupeodes corollae* (5.43%).

**Figure 9.**
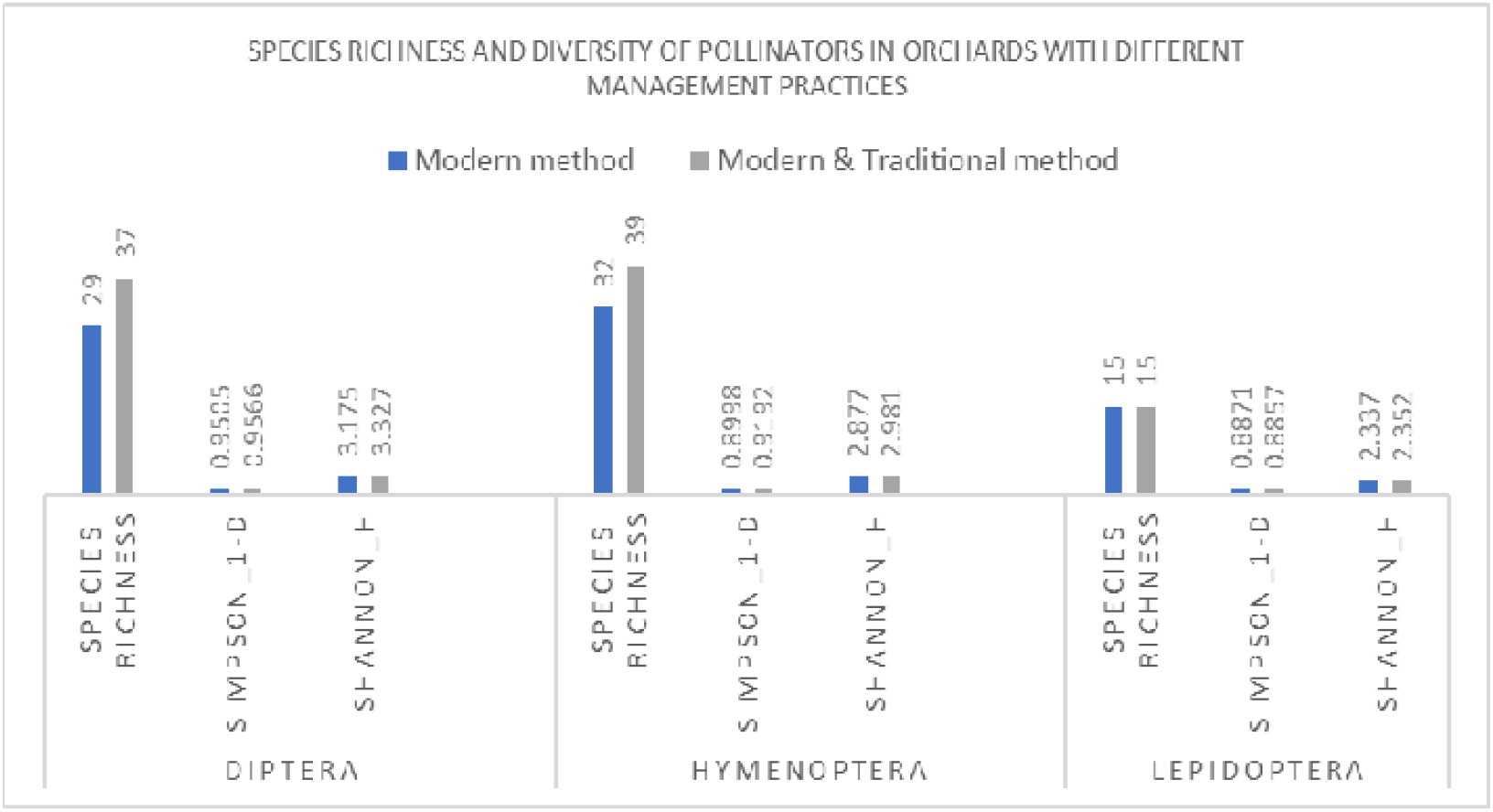
Account of species richness and diversity in orchards with different management practices

However, in orchards managed solely using modern techniques, the most abundant species in the Hymenoptera order were *Apis cerena indica* (26.5%), *Apis mellifera* (9.43%), *Megachilae lanata* (7.22%), and *Bombus haemorrhoidalis* (4.82%). For the Diptera order, most abundant species were *Meliscaeva cinctella* (9.67%), *Eristalis tenax* (8.54%), *Eupeodes corollae* (8.04%), *Chrysoma megacephala* (6.15%), *Episyrphys balteatus* (5.52%), and *Sphaerophoria indiana* (5.15%).

Regarding Lepidoptera, the most abundant species included *Aglais caschmirensis* (17.83% & 15.56%), *Colias fieldii fieldii* (16.43% & 15.56%), *Pieris brassicae* (11.42% & 18.99%), *Eurema hecabe* (10.32% & 9.84%), and *Pieris canidia* (11.42% & 6.17%) in orchards implementing both modern and traditional management practices and orchards with only modern management practices, respectively.

The entire Kullu valley is very much impacted by increased settlement and infrastructure. Orchards that are situated away from large settlement area exhibits higher pollinator diversity and richness in all three orders (Figure 10). Most abundant pollinators recorded in orchards situated far from large settlements included *Apis cerena indica* (23.68%), *Apis mellifera* (12.68%), *Megachilae lanata* (8.3%), *Osmia caerulescence* (6.69%), and *Bombus haemorrhoidalis* (4.9%) from the Hymenoptera order; *Eristalis ten*ax (9.01%), *Chrysoma megacephala* (7.32%), *Meliscaeva cinctella* (7.01%), *Eupeodes corollae* (5.92%), and *Episyrphys balteatus* (5.44%) from the order Diptera.

**Figure 10.**
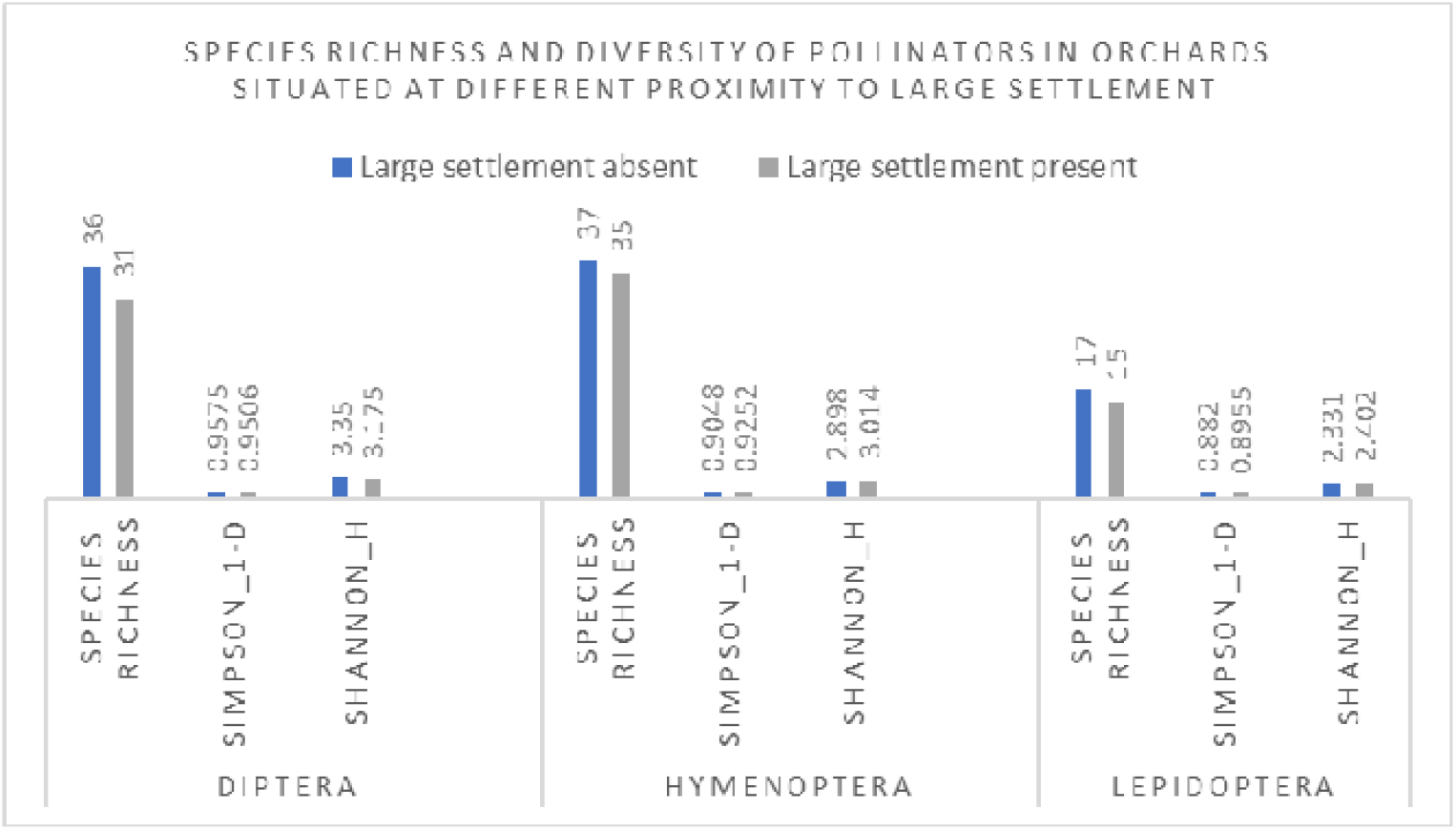
Account of species richness and diversity in orchards at different proximity to large settlement

On the other hand, and most abundant species in orchards located nearer to the settlement area include *Apis cerena indica* (18.97%), *Apis mellifera* (12.51%), *Megachilae lanata* (5.85%), Ceratina smaragdula (5.13%), Amegilla zonata (5.02%) and *Bombus haemorrhoidalis* (4.62%) from the order hymenoptera; *Eristalis tenax (8*.*14%), Episyrphys balteatus (7*.*35%), Eristalis horticola (7*.*17%), Chrysoma megacephala (7*.*17%), Eupeodes corollae (6*.*55%), Eristalis similis (6*.*28%), Sphaerophoria indiana (5*.*67%), and Eupeodes latifasciatus (5%) f*rom the order Diptera.

In case of lepidoptera most abundant species include, *Aglais caschmirensis* (16.89% & 17.45%), *Colias fieldii fieldii* (17.02% & 15.12%), *Pieris brassicae* (15.64% & 11.37%), *Eurema hecabe* (10.84% & 9.34%), and *Pieris canidia* (10.72% & 8.72%) in orchards away from large settlement and orchards near to large settlement respectively.

## Discussion

Present study suggested that hymenopteran and dipterans were the most effective pollinators with higher species diversity and richness followed by lepidoptera. Recent studies by Raj et al. (2012) and Mattu (2014) have also underscored the significance of hymenopterans and dipterans as the primary insect pollinators for apple and other temperate fruit crops in the Himalayan region. Apart from abundant species, some of the other important pollinators include solitary bees like *Xylocopa sp*., *Ceratina sp*., *Bombus sp*., *Amegilla sp*., *Osmia sp*., and other syrphid flies and drone flies. Wild bee species such as *Megachilae sp*., *Andrena sp*., *Xylocopa sp*., were found to be abundant in orchards which is less disturbed and situated near to the natural forest and far from large settlements and agricultural practices. This finding also supports the earlier study by Kremen et al. (2004) and more recent study by Mattu and Bhagat (2015). Additionally, it was observed that the diversity and relative abundance of majority of the insect pollinators were notably higher in orchards situated nearer to unmanaged and natural habitats and compared to those farther away from such habitats. These findings align with the conclusions drawn by Ricketts et al. (2008) and Joshi et al. (2016), where authors similarly found that both pollinator richness and visitation rates on crops decline with increasing distance from natural unmanaged habitats.

During the survey it was recorded that every farmer in this valley uses heavy number of pesticides on the crop for higher yield. In case of modern management farmers treat both the plant and soil with different pesticides, insecticides and fertilizers and clean the ground removing any wild growth along with other standard methods. On the other hand, where still management is done following traditional ways with adaptation to modern techniques, farmers use pesticides and insecticides along with organic fertilizers and doesn’t clean the sideways or old logs from the ground. Hence, it was found that species diversity was higher in orchards with both type of management interventions due to availability of natural resources for nesting and foraging as well. Species diversity and richness were comparatively less in orchards situated near agricultural lands. This can be due to the high use of pesticides in agricultural fields as well. This valley is also highly prone to natural and anthropogenic disasters. Apart from that tourisms and agricultural shifts are also prominent leading to frequent land use change. It was observed that in orchards near areas with large settlements and high tourism pressure were visited by a smaller number of wild pollinators. However, there are not much previous studies that show the impact of these type of land use practices on pollinators.

## Conclusion

The research highlights the rich diversity and abundance of pollinators across orchards within Kullu valley. Although there have been several studies on pollinators in Himachal Pradesh, including the Kullu district, few have investigated how diverse land use practices impact pollinator richness, diversity, and abundance on a landscape scale. Given the mosaic-like nature of the Kullu valley, a comprehensive approach of conservation and management involving all land use categories is crucial to sustain the natural habitats of pollinators in this region. Failure to do so might result in survival of only the more common and adaptable species in near future, while rarer species could face extinction risks. This study lays the basis for formulating sustainable management plans and strategies for pollinator conservation in this landscape. Additionally, it also suggests that conducting similar studies across different seasons and involving various crops could offer a more comprehensive understanding of the situation, facilitating improved management strategies.

Observations made during fieldwork also indicated that in orchards where commercial bee hives (Mainly *Apis mellifera*) were present, there was a noticeable decrease in the visitation rate of wild and native pollinators. However, detailed investigation needed to comprehend the impact of commercial bee hives on the visitation rate of wild and native pollinators in orchards of Kullu valley. This understanding could aid in devising more sustainable agricultural practices.

## Notes

### Competing Interest Statement

The authors have declared no competing interest.

## References

Ahmad F, Joshi SR, and Gurung MB. 2003. The Himalayan cliff bee Apis laboriosa and the honey hunters of the Himalayas. athmandu, Nepal: ICIMOD.

Aizen MA, and Feinsinger P. 1994. Forest fragmentation, pollination, and plant reproduction in a Chaco dry forest, Argentina. Ecology 75(2), pp.330–351.

Allen-Wardell G, Bernhardt P, Bitner R, Burquez A, Buchmann S, Cane J, Cox PA, Dalton V, Feinsinger P, Ingram M, and Inouye D. 1998. The potential consequence of pollinator declines on the conservation of biodiversity and stability of food crop yields. Conservation biology pp.8–17.

Arora GS. 1990. Collection and preservation of animals (Lepidoptera), Zoological Survey of India, Calcutta, 131–138.

Arya MK, and Badoni A. 2023. Insect Pollinator Assemblage on Temperate Fruits Crops in Kumaun Himalaya. Indian Journal of Entomology pp.315–321.

Biesmeijer JC, Roberts SP, Reemer M, Ohlemüller R, Edwards M, Peeters T, Schaffers AP, Potts SG, Kleukers R, Thomas CD, and Settele J. 2006. Parallel declines in pollinators and insect-pollinated plants in Britain and the Netherlands. Science 313(5785), pp.351–354.

Cameron SA, Lozier JD, Strange JP, Koch JB, Cordes N, Solter LF, and Griswold TL. 2011. Patterns of widespread decline in North American bumble bees. Proceedings of the National Academy of Sciences 108(2), pp.662–667.

Chauhan M, Uniyal VP, Chandra A, Thakur P, and Mehrwar V. 2021. Preliminary assessment and conservation of insect pollinators through community participation in the Lahaul and Spiti district of Himachal Pradesh, India. Current Science 120(5), p.883.

Dixon P. 2003. VEGAN, a package of R functions for community ecology. Journal of vegetation science, 14(6), pp.927–930.

Gallai N, Salles JM, Settele J, and Vaissière BE. 2009. Economic valuation of the vulnerability of world agriculture confronted with pollinator decline. Ecological economics 68(3), pp.810–821.

Gardner J, Sinclair J, Berkes F, and Singh R. 2002. Accelerated Tourism Development and Its Impacts in Kullu-Manali, H.P., India. Tourism Recreation Research, 27, pp. 20–9. 10.1080/02508281.2002.11081370.

Ghazoul J. 2005. Buzziness as usual? Questioning the global pollination crisis. Trends in ecology & evolution 20(7), pp.367–373.

Gibson RH, Knott B, Eberlein T, and Memmott J. 2011. Sampling method influences the structure of plant-pollinator. Oikos 120: 822–831. 10.1111/j.1600-0706.2010.18927.x

District Wise Disaster Vulnerability of the State. 2018. Available at: https://hpsdma.nic.in/index1.aspx?lsid=72&lev=2&lid=14&langid=1 Himachal Pradesh State disaster management authority [Date accessed: 02 September 2023].

IPBES. 2016. The assessment report of the Intergovernmental Science-Policy Platform on Biodiversity and Ecosystem Services on pollinators, pollination and food production. S.G. Potts, V. L. Imperatriz-Fonseca, and H. T. Ngo, (eds). Secretariat of the Intergovernmental Science-Policy Platform on Biodiversity and Ecosystem Services, Bonn, Germany. 552 pages.

Joseph ANT. 1990. Collection and preservation of animals (Diptera), Zoological Survey of India, Calcutta, 141–144.

Joshi N, Otieno M, Rajotte E, Fleischer S, and Biddinger D. 2016. Proximity to Woodland and Landscape Structure Drives Pollinator Visitation in Apple Orchard Ecosystem. Frontiers in Ecology and Evolution 4. 10.3389/fevo.2016.00038.

Kapkoti B, Joshi RK, and Rawal RS, 2016. Variations in the abundance and diversity of insects in apple orchards of Kumaun, Western Himalaya, India. Current Science, pp.438–443.

Kennedy CM, Lonsdorf E, Neel MC, Williams NM, Ricketts TH, Winfree R, Bommarco R, Brittain C, Burley AL, Cariveau D, and Carvalheiro LG. 2013. A global quantitative synthesis of local and landscape effects on wild bee pollinators in agroecosystems. Ecology letters 16(5), pp.584–599.

Kindt R. 2019. Package ‘BiodiversityR’. Package for community ecology and suitability analysis, 2, pp.11–12.

Klein AM, Vaissiere BE, Cane JH, Steffan-Dewenter I, Cunningham SA, Kremen C, and Tscharntke T. 2007. Importance of pollinators in changing landscapes for world crops. Proceedings of the royal society B: biological sciences 274(1608), pp.303–313.

Kremen C, William NM, Bugg RL, Fay JP, and Thorp RW. 2004. The area requirements of an ecosystem service: crop pollination by native bee communities in California, Ecology Letters 7: 1109–19.

Mattu VK. 2014. Role of honeybees and other pollinators in crop productivity and impacts of climate change. Workshop on Promotion of Honey Beekeeping in Haryana, Panchkula, pp. 56–74.

Mattu VK, and Bhagat T. 2015. Effect of changing landscapes on diversity, distribution and relative abundance of insect pollinators on apple crop in Northwest Himalayas. International Journal of Science & Research, 4(5), pp.1762–1767.

Misganaw M, Mengesha G, and Awas T. 2017. Perception of farmers on importance of insect pollinators in Gozamin district of Amhara region, Ethiopia. Biodiversity International Journal 1(5), pp.54–60. 10.15406/BIJ.2017.01.00029.

National Research Council, 2007. Status of pollinators in North America. National Academies Press.

Negi V, and Irfan M, 2022. Land Use/Cover Mapping and Change Detection Using Remote Sensing Techniques: A Case of Upper Kullu Valley, Himachal Pradesh. Current World Environment 17(2), pp.417–426.

Oldroyd BP. 2007. What’s killing American honeybees? PLoS Biology 5(6), p.e168.

Orford K, Vaughan I, and Memmott J. 2015. The forgotten flies: the importance of non-syrphid Diptera as pollinators. Proceedings of the Royal Society B: Biological Sciences 282. 10.1098/rspb.2014.2934.

Partap T, and Partap U. 2001. Declining apple production and worried Himalayan farmers: promotion of honeybees for pollination.

Partap U. 2010a. Honeybees and ecosystem services in the Himalayas. In Spehn, EM; Rudmann-Maurer, K. Körner, C; Maselli, D (eds), Mountain biodiversity and global change, pp. 21–21. Basel, Switzerland: Global Mountain Biodiversity Assessment (GMBA) of DIVERSITAS.

Partap U. 2010b. Innovations in revival strategies for declining pollinators with particular reference to the indigenous honeybees: Experiences of ICIMODLs initiatives in the Hindu Kush-Himalayan region. Pest Management and Economic Zoology 18, pp.85–95.

Partap U, and Partap, T. 1997. Managed crop pollination: the missing dimension of mountain productivity. International Centre for Integrated Mountain Development (ICIMOD): Discussion paper series.

Partap U, and Partap T. 2002. Warning signals from apple valleys of the HKH region: pollination problems and farmers’ management efforts. Kathmandu, Nepal: International Centre for Integrated Mountain Development (ICIMOD).

Potts SG, Biesmeijer JC, Kremen C, Neumann P, Schweiger O, and Kunin WE. 2010. Global pollinator declines: trends, impacts and drivers. Trends in ecology & evolution 25(6), pp.345–353.

Rader R, Bartomeus I, Garibaldi LA, Garratt MP, Howlett BG, Winfree R, Cunningham SA, Mayfield MM, Arthur AD, Andersson GK, and Bommarco R. 2016. Non-bee insects are important contributors to global crop pollination. Proceedings of the National Academy of Sciences 113(1), pp.146–151.

Rader R, Cunningham S, Howlett B, and Inouye D. 2020. Non-Bee Insects as Visitors and Pollinators of Crops: Biology, Ecology and Management. Annual review of entomology 10.1146/annurev-ento-011019-025055.

Raj H, Mattu VK, and Thakur ML. 2012. Pollinator diversity and relative abundance of insect visitors on apple crop in Shimla hills of western Himalaya, India. International Journal of Science and Nature 3 (3): 507–513.

Ricketts TH, Regetz J, Dewenter IS, Cunningham SA, Kremen C, Bogdanski A, Herren BG, Greenleaf SS, Klein AM, Mayfield MM, Morandin LA, Ochieng A, and Viana BF. 2008. Landscape effects on crop pollination services: are there general patterns? Ecological Letters 11: 499–515.

Sharma H, and Chauhan S. 2013. Agricultural Transformation in Trans Himalayan Region of Himachal Pradesh: Cropping Pattern, Technology Adoption and Emerging Challenges. Agricultural Economics Research Review 26, pp. 173–179.

Sharma HK, Bakshi N, Thakur RK, and Devi M. 2016. Diversity and density of insect pollinators on sweet cherry (Prunus avium L.) in temperate region of Kullu valley of Himachal Pradesh. Journal of Entomological Research 40(2), pp.123–128.

Sharma HK, Gupta JK, Suman V, and Belavadi VV. 2015. Studies on flowering phenology, pollinator diversity, supplemented pollination and their impact on fruit set in apple under changing climatic scenario in Kullu valley of Himachal Pradesh. International Journal of Farm Sciences, 5(4), pp.156–164.

Sharma LK, and Rana RK. 2015. Effect of top working to improve pollination in apple (Malus x domestica Borkh.) orchards under mid hill conditions of Kullu district. Himachal Journal of Agricultural Research 41(1), pp.89–92.

Sharma, R. and Gupta, J.K., 2010. Effect of changing landscapes on the density and diversity of insect pollinators on apple crop in Kullu valley of Himachal Pradesh. Pest Management and Economic Zoology, 18(1/2), pp.277–280.

Sharma RM and Mitra B. 2011. A preliminary study on insect pollinators of temperate fruit crops in Himachal Pradesh. Records of the Zoological Survey of India 111(3), pp.103–110.

Slariya M. 2014. Shifting from Traditional Food Cropping to Cash Cropping pp. 83–92. 10.1007/978-4-431-54868-3_7.

Steffan-Dewenter, I, Potts SG, and Packer L. 2005. Pollinator diversity and crop pollination services are at risk. Trends in ecology & evolution 20(12), pp.651–652.

Thakur B, and Mattu VK. 2014. Diversity and distribution of pollinators of temperate fruit crops of Shimla hills in Himachal Pradesh. Asian Journal of Advanced Basic Sciences 3(1), pp.151–163.

Ulyshen M, UrbanLMead K, Dorey J, and Rivers J. 2023. Forests are critically important to global pollinator diversity and enhance pollination in adjacent crops. Biological reviews of the Cambridge Philosophical Society 10.1111/brv.12947.

Vaidya P, Bhardwaj SK, and Sood S. 2018. Land use and land cover changes in Kullu valley of Himachal Pradesh. Indian Journal of Agricultural Sciences 88(6), pp.902–906.

Vanbergen AJ, and Initiative TIP. 2013. Threats to an ecosystem service: pressures on pollinators. Frontiers in Ecology and the Environment 11(5), pp.251–259.

Verma LR, and Partap U. 1993. The Asian hive bee, Apis cerena, as a pollinator in vegetable seed production. International Centre for Integrated Mountain Development (ICIMOD).

Verma MK, Ahmed N, Singh AK, and Awasthi OP. 2010. Temperate tree fruits and nuts in India. Chronica Horticulture 50(4), pp.43–48.

Viana BF, Boscolo D, Mariano Neto E, Lopes LE, Lopes AV, Ferreira PA, Pigozzo CM, and Primo LM. 2012. How well do we understand landscape effects on pollinators and pollination services? Journal of Pollination Ecology 7.

Westphal C, Bommarco R, Carré G, Lamborn E, Morison N, Petanidou T, Potts SG, Roberts SP, Szentgyörgyi H, Tscheulin T, and Vaissière BE. 2008. Measuring bee diversity in different European habitats and biogeographical regions. Ecological monographs, 78(4), pp.653–671.

Williams PH, and Osborne JL. 2009. Bumblebee vulnerability and conservation world-wide. Apidologie, 40(3), pp.367–387.

